# Chromatin accessibility changes between *Arabidopsis* stem cells and mesophyll cells illuminate cell type-specific transcription factor networks

**DOI:** 10.1101/213900

**Authors:** Paja Sijacic, Marko Bajic, Elizabeth C. McKinney, Richard B. Meagher, Roger B. Deal

## Abstract

**Background:** Cell differentiation is driven by changes in transcription factor (TF) activity and subsequent alterations in transcription. To study this process, differences in TF binding between cell types can be deduced by methods that probe chromatin accessibility. We used cell type-specific nuclei purification followed by the Assay for Transposase Accessible Chromatin (ATAC-seq) to delineate differences in chromatin accessibility and TF regulatory networks between stem cells of the shoot apical meristem (SAM) and differentiated leaf mesophyll cells of *Arabidopsis thaliana.*

**Results:** Chromatin accessibility profiles of SAM stem cells and leaf mesophyll cells were highly similar at a qualitative level, yet thousands of regions of quantitatively different chromatin accessibility were also identified. We found that chromatin regions preferentially accessible in mesophyll cells tended to also be substantially accessible in the stem cells as compared to the genome-wide average, whereas the converse was not true. Analysis of genomic regions preferentially accessible in each cell type identified hundreds of overrepresented TF binding motifs, highlighting a set of TFs that are likely important for each cell type. Among these, we found evidence for extensive co-regulation of target genes by multiple TFs that are preferentially expressed in one cell type or the other. For example, a set of zinc-finger TFs appear to control a suite of growth-and development-related genes specifically in stem cells, while another TF set co-regulates genes involved in light responses and photosynthesis specifically in mesophyll cells. Interestingly, the TFs within both of these sets also show evidence of extensively co-regulating each other.

**Conclusions:** Quantitative analysis of chromatin accessibility differences between stem cells and differentiated mesophyll cells allowed us to identify TF regulatory networks and downstream target genes that are likely to be functionally important in each cell type. Our findings that mesophyll cell-enriched accessible sites tend to already be substantially accessible in stem cells, but not vice versa, suggests that widespread regulatory element accessibility may be important for the developmental plasticity of stem cells. This work also demonstrates the utility of cell type-specific chromatin accessibility profiling in quickly developing testable models of regulatory control differences between cell types.

## Background

In higher plants, all above ground tissues are continuously produced due to the activities of self-renewing, pluripotent stem cells located in the central zone of the shoot apical meristem (SAM). Upon stem cell division, a subset of daughter cells is gradually displaced to the peripheral zones of the SAM where these cells continue to divide and differentiate. During this process, differentiating cells undergo transcriptional reprogramming as they acquire specialized fates within developing leaf primordia at the flanks of the SAM [1–2].

Chromatin compaction within the nucleus often restricts the access of transcription factors (TFs) to cis-regulatory elements, such as promoters and enhancers [3]. During differentiation, cells employ various mechanisms to induce local changes in chromatin properties, thereby modifying the accessibility of regulatory chromatin regions to the transcriptional machinery [3–4]. This ultimately leads to the establishment of lineage-specific TF regulatory modules and the resulting transcriptional output characteristic of a given cell type. To date, a limited number of such cell type-specific TF regulatory modules have been identified in plants. One well-studied example is the regulatory network of TFs that controls specification of the root non-hair cell type in the *Arabidopsis* root epidermis. In this system, the interactions of multiple TFs dictate expression of the non-hair fate master regulator, GLABRA2 (GL2), which subsequently determines cell fate [5–6]. This complex of TFs that regulate the expression of *GL2* was delineated through extensive genetic studies in numerous laboratories and now represents one of the best understood fate specification pathways in plants. To expedite mechanistic studies of cell fate specification in many other cell types, it will be important to be able to identify cell-type specific cis-regulatory regions and the transcription factors that act on them.

To measure DNA accessibility and TF binding, genome-wide analysis methods such as DNase I treatment of nuclei coupled with high-throughput sequencing (DNase-seq) have been used [7–9]. Mapping of DNase I hypersensitive sites (DHSs) allows for the identification of cis-regulatory elements because DHSs represent open chromatin regions where protein binding to DNA has displaced nucleosomes, generating a nuclease sensitive zone [10]. Large-scale DNase-seq studies have been instrumental in identifying cell type-specific *cis*-regulatory elements, most notably including a study involving more than 100 human cell types [11]. One substantial drawback of this powerful technique, however, is the requirement for large quantities of nuclei as starting material. Recently, the simple and sensitive Assay for Transposase-Accessible Chromatin with high-throughput sequencing (ATAC-seq) has been described [12–13], which requires much smaller amount of input material (~500 to 50,000 nuclei) [14]. In this method, a hyperactive Tn5 transposase loaded with sequencing adapters acts to simultaneously fragment and tag a genome with these adapters. Mapping of the transposase hypersensitive sites allows for detection of highly accessible chromatin regions and subsequent identification of TF binding sites within these regions [14].

One of the main limitations to successfully identifying cell type-specific *cis-*regulatory regions and studying the networks of transcription factors that bind to these elements is the difficulty in isolating specific cell types. The INTACT (Isolation of Nuclei TAgged in specific Cell Types) technique is one solution to this problem that is highly amenable to chromatin studies [15–16]. This system utilizes transgenic plants carrying two transgenes. The first encodes the nuclear targeting fusion (NTF) protein, which is comprised of a nuclear envelope-targeting domain, green fluorescent protein (GFP), and biotin ligase recognition peptide (BLRP). The second transgene is the *E. coli* biotin ligase (BirA) which specifically biotinylates the NTF protein. The *BirA* transgene is expressed from a constitutively active promoter, while the expression of NTF is driven by a cell type-specific promoter. The co-expression of these transgenes results in the biotinylation of nuclei in a specific cell type, which can then be affinity purified with streptavidin-coated magnetic beads.

In this study, we employed INTACT and ATAC-seq methods, collectively called INTACT-ATAC-seq, to identify and compare accessible chromatin regions between two distinct plant cell types: pluripotent stem cells in the central zone of the SAM, and highly specialized, fully-differentiated leaf mesophyll cells that originate from the stem cells of the SAM. The comparison of these two cell types offers a unique insight into chromatin dynamics and transcriptional regulatory control at both the starting and ending points of the differentiation process. Our results show that while most Transposase Hypersensitive Sites (THSs) are shared between both cell types, thousands of regions could be identified that were quantitatively more accessible in one cell type compared to the other. Furthermore, we identified transcription factor (TF) binding motifs within these THSs and used this information, in combination with publicly available expression and protein interaction data, to build cell-specific TF-to-TF regulatory networks, and to predict the downstream target genes of these TF networks. Our results suggest that distinct classes of TFs collaborate to produce cell type-specific transcriptomes in the stem cell and mesophyll cell types. We also demonstrate that INTACT-ATAC-seq is a powerful technique to quickly develop testable hypotheses regarding TF regulatory networks and their roles in cell fate specification.

## Results

### Validation of cell type-specific INTACT lines and INTACT-ATAC-seq data

The *CLAVATA3 (CLV3)* gene, a known stem cell marker [17], is exclusively expressed in the meristematic stem cells found in the central zone of the SAM [18]. We used the upstream and downstream regulatory sequences of *CLV3* to drive the expression of the nuclear targeting fusion (NTF) transgene selectively in the SAM stem cells. Expression of the *CLV3p::NTF* construct in *CLV3p::NTF;ACT2p::BirA T*_2_ transgenic plants was confirmed using confocal microscopy by visualizing the Green Fluorescent Protein (GFP), which is a part of the NTF, specifically in the central zone of the SAM (Figure 1A). Similarly, the promoter of the *Rubisco small subunit 2B (RBC)* gene, active only in the mesophyll cells [19], was used to drive the expression of the NTF in leaf mesophyll cells. The expression of this construct was visualized by confocal microscopy in the leaves of the *RBCp::NTF;ACT2p::BirA T*_*2*_ transgenic plants. GFP expression was observed in the inner cell layers of the sectioned leaf, and is excluded from the leaf epidermis (Figure 1A).

**Figure 1.**
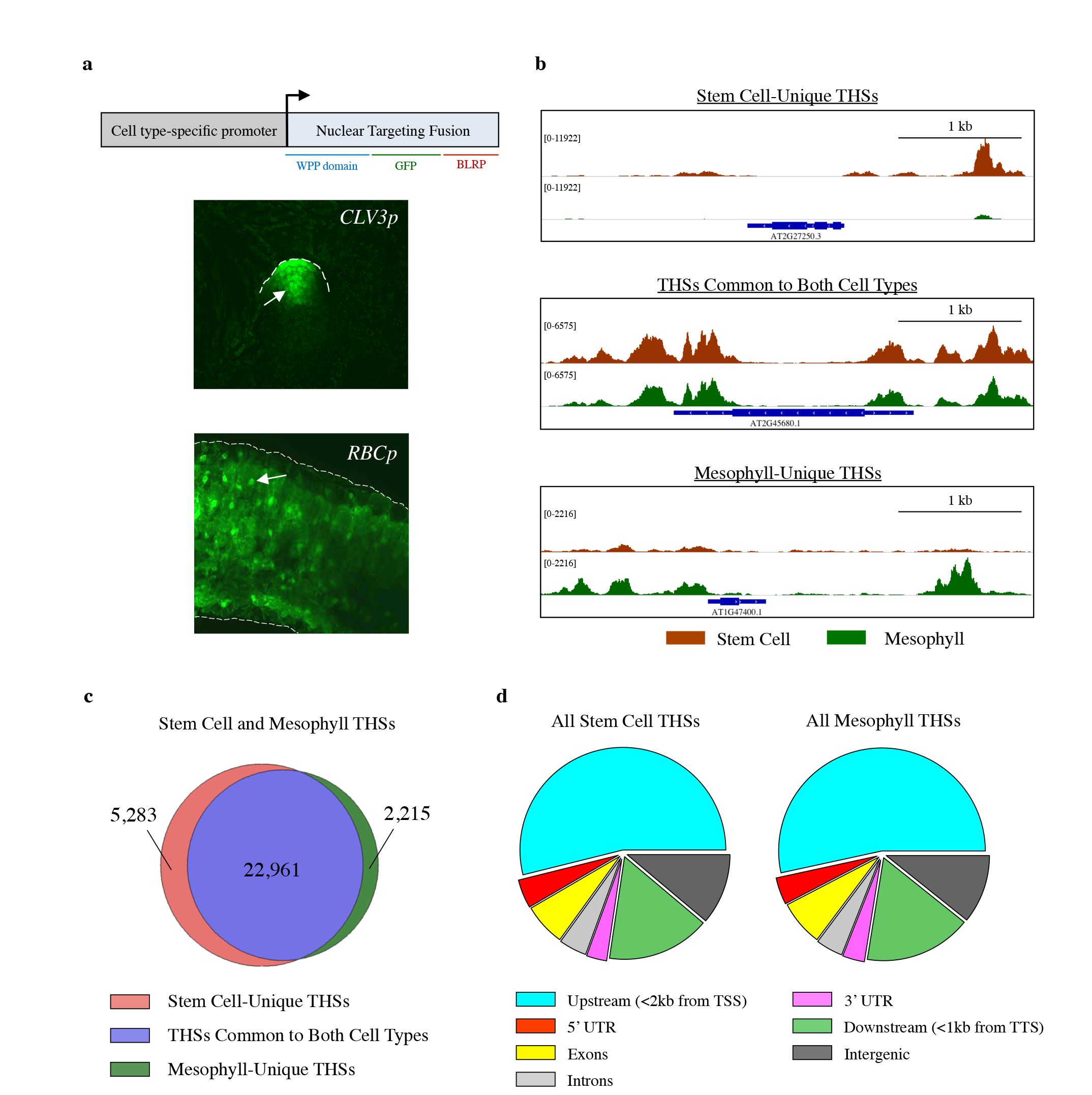
Characterization of INTACT transgenic lines and overview of ATAC-seq data from each cell type. **(a)** The upper panel is a schematic representation of the Isolation of Nuclei TAgged in specific Cell Types (INTACT) system for isolating nuclei from specific cell types. The Nuclear Targeting Fusion (NTF) contains a WPP nuclear envelope-binding domain, Green Fluorescent Protein (GFP) for visualization, and a biotin ligase recognition peptide (BFRP), which can be biotinylated by the BirA biotin ligase. BirA is expressed constitutively while NTF is driven from a cell type specific promoter. When these transgenes are coexpressed in a cell the nucleus becomes biotinylated, allowing all nuclei of that cell type to be selectively purified with streptavidin beads. Below the gene diagram are confocal images of GFP expression in the ***CLV3p:NTF;ACT2p:BirA*** line (upper) and ***RBCp:NTF;ACT2p:BirA*** line (lower), showing NTF expression in the shoot apical meristem and mesophyll cells, respectively. Fluorescent nuclei are labeled with arrowheads, **(b)** Three Integrated Genome Viewer (IGV) snapshots of normalized ATAC-seq reads from shoot apical stem cell (red) and mesophyll (green) nuclei. Different categories of Transposase Hypersensitive Sites (THSs) are observed: Top panel) Stem cell-unique: THSs identified only in stem cells; Middle panel) Common to both cell types: THSs that were identified in both stem cells and mesophyll cells; and Bottom panel) Mesophy11-unique: THSs that were identified only in mesophyll cells. **(c)** Overlap of stem cell and mesophyll ATAC-seq THSs identified by peak calling in at least two biological replicates of that cell type. **(d)** Genomic distribution of all the THSs identified in two replicates for either stem cell or mesophyll ATAC-seq.

The INTACT protocol for cell type-specific nuclei purification from *CLV3p::NTF;ACT2p::BirA* and *RBCp::NTF;ACT2p::BirA T*_2_ transgenic plants was performed as previously described [20]. A total of 25,000 freshly isolated nuclei were used for ATAC-seq, and three biological replicates were performed per cell type. In parallel, we performed ATAC-seq on genomic DNA isolated from leaf tissue as a control for sequence-specific Tn5 transposase incorporation bias. More than 84 million reads were obtained for each biological replicate through paired-end sequencing (Figure S1A). After aligning the ATAC-seq reads to the *Arabidopsis thaliana* TAIR10 genome, we found that, on average, 46% of all reads from the stem cells and 21% from the mesophyll cells were successfully mapped to the nuclear genome, with the remainder of reads mapping to organelle genomes (Figure S1A). This level of organelle DNA carry over was unexpected based on our previous INTACT purifications from root tissue, and is likely attributable to the sheer abundance of chloroplasts in shoot tissue. All reads that aligned to the organellar genomes were subsequently omitted from downstream analyses. More than 15 million reads per replicate passed the quality filtering stage of analysis (Figure S1A), which is more than sufficient to successfully identify accessible chromatin regions in *Arabidopsis*, as has been recently demonstrated [14]. The processed, alignment files were compared using principal component analysis (PCA) [21]. The six libraries segregated by cell type, with low variation between replicates, indicating the high level of reproducibility in our datasets (Figure S1B). For each library, we analyzed the fragment size distribution of the aligned reads to determine the number of nucleosome-containing (>150 bp) and nucleosome-free reads (<150 bp). Nucleosome-free reads are regions of accessible chromatin where a transcription factor is likely bound. Conversely, nucleosome-containing reads are less accessible to transcription factor binding and are therefore less relevant to the scope of this study. In the stem cell and mesophyll ATAC-seq datasets, we saw a fragment size distribution primarily of 100 bp fragments and smaller, indicating that our libraries were composed of primarily nucleosome-free reads (Figure S1C). Additionally, the periodic dips in the size distribution graphs demonstrate a clear pattern of the helical pitch of DNA, further confirming that our transposase treatment was of sufficient coverage. The fragment size distribution for genomic DNA ATAC-seq library was smaller, primarily 50 bp in size, and lacked a clear representation of the helical pitch of DNA (Figure S1C). In summary, INTACT-ATAC-seq is a very effective method for obtaining a large quantity of highly reproducible accessible chromatin reads in *Arabidopsis* mesophyll and stem cells.

### Identification and genomic distribution of cell type-specific accessible chromatin regions

Since the ATAC-seq data among all replicates were highly reproducible (Figure S1B), we focused our analysis on the two biological replicates with the highest number of aligned reads for each cell type. To keep our analysis consistent across samples, we first scaled the reads from each cell type to the same sequencing depth (15,288,699 reads, Figure S1A) and then used the peak calling function of the HOMER package [22] to identify open chromatin regions. From this set of transposase hypersensitive sites (THSs) identified by HOMER, we examined only the THS regions that were identified in both replicates of each cell type, which we refer to as reproducible THSs (Table S1). The majority of these reproducible THSs (22,961 of 30,459 sites) were common to both cell types, while 5,283 and 2,215 THSs were reproducibly called only in one cell type (stem cells and mesophyll cells, respectively) (Figure 1B and C). The genomic distribution of reproducible THSs is very similar between the two cell types, with 53% of THSs located within 2 kb upstream of the gene transcription start sites (TSSs), 18% located within the gene body, 16% located within 1 kb downstream of transcription termination sites (TTSs), and 10% located in the intergenic region (Figure 1D). This genomic distribution of reproducible THSs suggests that the majority of cis-regulatory regions in *Arabidopsis* genome are located in the vicinity of gene core promoters, as previously observed in other *Arabidopsis* cell types [23].

Since the majority of identified THSs were common to both cell types (Figure 1C) we hypothesized that there may still be quantitative differences between cell types at the shared THSs that would not be identified by our all-or-nothing peak calling approach. To examine quantitative differences in accessible chromatin regions between the two cell types, we calculated the normalized total read counts at each THS in the merged set of reproducible THSs for both cell types *(i.e.* all THSs shown in Figure 1C). The calculated read counts were then evaluated using DESeq2 to identify reproducible quantitative differences in accessibility between cell types [24] (Table S2). Only those THSs that had a log fold change of 1 or more in a specific cell type were categorized as THSs enriched in that cell type (see Methods).

With this approach we identified a total of 7,394 THSs that are more accessible in stem cells and 5,895 THSs that are more accessible in mesophyll cells (Figure 2A). This analysis captured the majority of THSs originally identified as cell type-unique by peak calling, and added several thousand differential THSs to each cell type that were previously classified as being present in both cell types by peak calling alone. We now refer to these collections of THSs that are quantitatively significantly different between cell types as cell type-enriched THSs.

**Figure 2.**
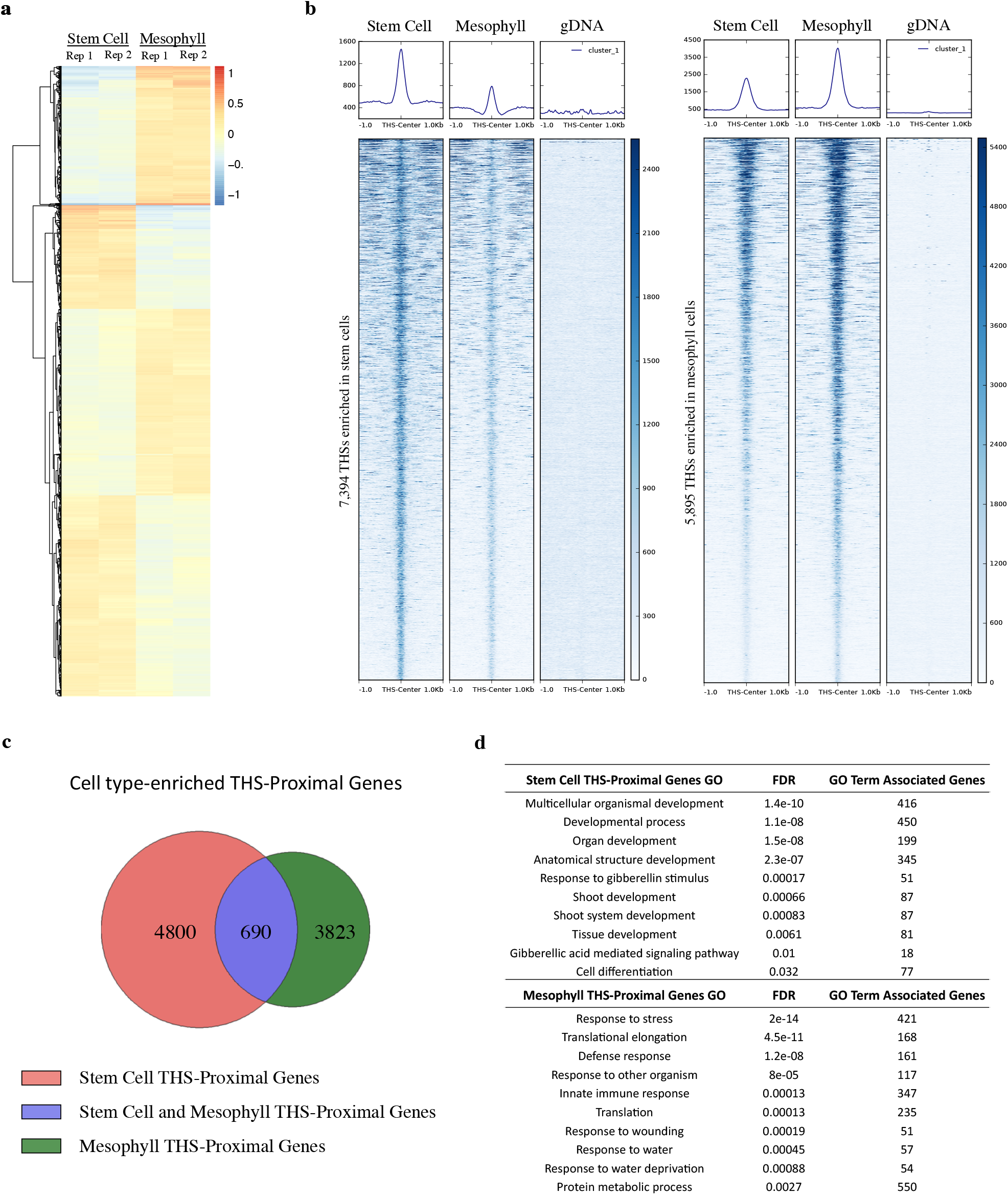
Chromatin accessibility differences between stem cells and mesophyll cells. **(a)** Heatmap showing the log ratio of normalized read count of the top 13,289 THSs that are statistically different between stem cell and mesophyll ATAC-seq samples. Each line on the heatmap represents a single THS, and the values at that region are given for each of two replicates in each cell type. Increased chromatin accessibility between the four samples is colored red and decreased chromatin accessibility is colored blue, compared to an average value set to 0. **(b)** Normalized read signal in stem cell, mesophyll, and genomic DNA ATAC-seq samples over cell type-enriched THS regions. The left set of panels show ATAC-seq signal over the 7394 stem cell-enriched THSs, while the right set of panels shows ATAC-seq signal over the 5,895 THSs enriched in mesophyll cells. **(c)** Each cell type-enriched THS was assigned to its nearest TSS as the putatively regulated target gene. Venn diagram shows the overlap of cell type-enriched THS-proximal genes. **(d)** Examples of 10 GO terms that were found only among the lists of genes that have a nearby cell type-enriched THS in a given cell type (i.e. from the non-overlapping portions of the diagram in (c)). FDR = False Discovery Rate, GO = Gene Ontology.

Each set of cell type-enriched THSs had a similar genomic distribution which matched the trend of the overall THS distribution, with more than 75% of these THSs mapping within 2 kb upstream of the gene TSSs and 1 kb downstream of TTSs (Figure S2A). Heatmaps and average plots of the ATAC-seq signal from mesophyll, stem cell, and genomic DNA datasets were visualized over stem cell-enriched and mesophyll-enriched THSs to examine the chromatin accessibility differences between cell types at these sites (Figure 2B). At stem cell-enriched THSs, the stem cell ATAC-seq signal is strongest in the centers of these regions (with an average maximum of approximately 1500 RPKM) and drops off sharply to either side. In contrast, the mesophyll cells show far less accessibility in these regions, but are nonetheless accessible to transposase integration to some degree (Figure 2B, left panels). At mesophyll-enriched THSs, the mesophyll cells show the highest accessibility in these regions with an average maximum of 4,400 RPKM (Figure 2B, right panels). It is worth noting that this read signal is much higher than that seen in stem cells at stem cell-enriched THSs. Interestingly, stem cells also show relatively high accessibility at mesophyll-enriched THSs, with an average maximum in the same range as that seen at stem cell-enriched THSs. These results strongly indicate that mesophyll-enriched THSs are already highly accessible in stem cells, but not vice versa.

On the other hand, ATAC-seq reads from genomic DNA were present at negligible levels at both the stem cell-enriched and mesophyll-enriched THSs (Figure 2B). In fact, we identified only 35 THSs in genomic DNA by peak calling, with approximately 75% of them located in the intergenic regions of the genome (Figure S3A and B). These results suggest a very low level of Tn5 integration bias at this scale. Taken together, we successfully used INTACT-ATAC-seq to identify cell type-enriched THSs, which reflect the reproducible differences in the chromatin accessibility between the stem cells and mesophyll cells.

### Gene ontology analysis of the genes associated with cell type-enriched THSs

THSs represent accessible, nucleosome-free, chromatin regions and are likely to contain cis-regulatory elements that control the expression of nearby genes. To identify the genes associated with the cell type-enriched THSs we used the PeakAnnotator program [25] to assign each THS to the nearest gene TSS, regardless of whether the TSS is upstream or downstream. We will hereafter refer to the genes associated with the stem cell-enriched THSs as the stem cell THS-proximal genes, and the genes associated with the mesophyll-enriched THSs as the mesophyll THS-proximal genes. The 7,394 stem cell-enriched THSs mapped to the 5,490 stem cell THS-proximal genes, while the 5,895 mesophyll-enriched THSs mapped to the 4,513 mesophyll THS-proximal genes (Figure 2C and Figure S2B). These results indicate that in each cell type a single gene sometimes has more than one cell type-enriched THS associated with it, while the majority of genes that have a nearby cell type-enriched THS are associated with a single such site. As shown in Figure S2B, a greater number of ATAC-seq reads were observed across the gene bodies of these THS-proximal gene sets for the cell type they were originally identified in. In other words, the stem cell THS-proximal genes showed more ATAC-seq reads across their gene bodies in the stem cell dataset compared to the mesophyll dataset, and vice versa (Figure S2B). These results suggest that the proximal genes of enriched THSs have more accessible chromatin across their gene body, and therefore are more likely to be highly transcribed in the cell type where the THS is enriched. It was also found that the majority of ATAC-seq reads relative to these genes are localized proximally upstream of the transcription start sites (TSSs) and downstream of the transcript end site (TES) (Figure S2B). Minimal transposase bias was found in these analyses, and was primarily confined to gene bodies in both sets of genes. Importantly, however, such bias was not observed at the specific sites where the majority of our enriched THSs were located (Figure 2B, S2B).

We next used AgriGO [26–27] to identify overrepresented Gene Ontology (GO) terms within the THS-proximal gene sets for each cell type (Table S3). We focused our analysis only on the GO terms that had a false discovery rate (FDR) of less than 0.05. This analysis revealed that many of the genes associated with the stem cell-enriched THSs are involved in the regulation of cell differentiation and shoot development, while the genes proximal to the mesophyll-enriched THSs were predominantly involved in response to biotic and abiotic stimuli, which is consistent with the known functions of these two cell types (Figure 2D).

### Enriched motif analysis and identification of cell type-specific transcriptional regulatory networks

As described above, THSs represent more accessible chromatin regions, which likely contain TF binding sites that can recruit TFs to regulate the expression of nearby genes. To identify specific transcription factors that may play important roles in establishing and maintaining the stem cell and mesophyll cell fates during development, we first identified sequence motifs that were overrepresented in cell type-enriched THSs. This was achieved by performing a MEME-ChIP analysis on the repeat-masked sequences within these THS regions [28]. We discovered a total of 316 overrepresented motifs within the stem cell-enriched THSs and 211 motifs within mesophyll-enriched THSs (Figure 3A and Table S4). Next, to determine which TFs show differential expression in one cell type or other, we ranked the TFs that bind the identified motifs based on their expression level in each cell type using publicly available RNA-seq and microarray data [29–30]. This was done by first calculating the relative expression rank by percentile for each gene in these datasets (see Methods). Then, the difference in expression rank for each TF of interest was measured between the two cell types (Table S5). In total, we identified 23 stem cell-enriched and 129 mesophyll-enriched TFs that have at least a two fold difference in their relative expression ranking between cell types (Figure 3 and Table S5). We then used these TF sets as input for the STRING database, which combines both known protein-protein interactions and functional interactions among genes (*e.g.* co-expression, text mining association, interactions in orthologs from other species, etc.) to infer and predict functional connections between a set of input genes [31].

**Figure 3.**
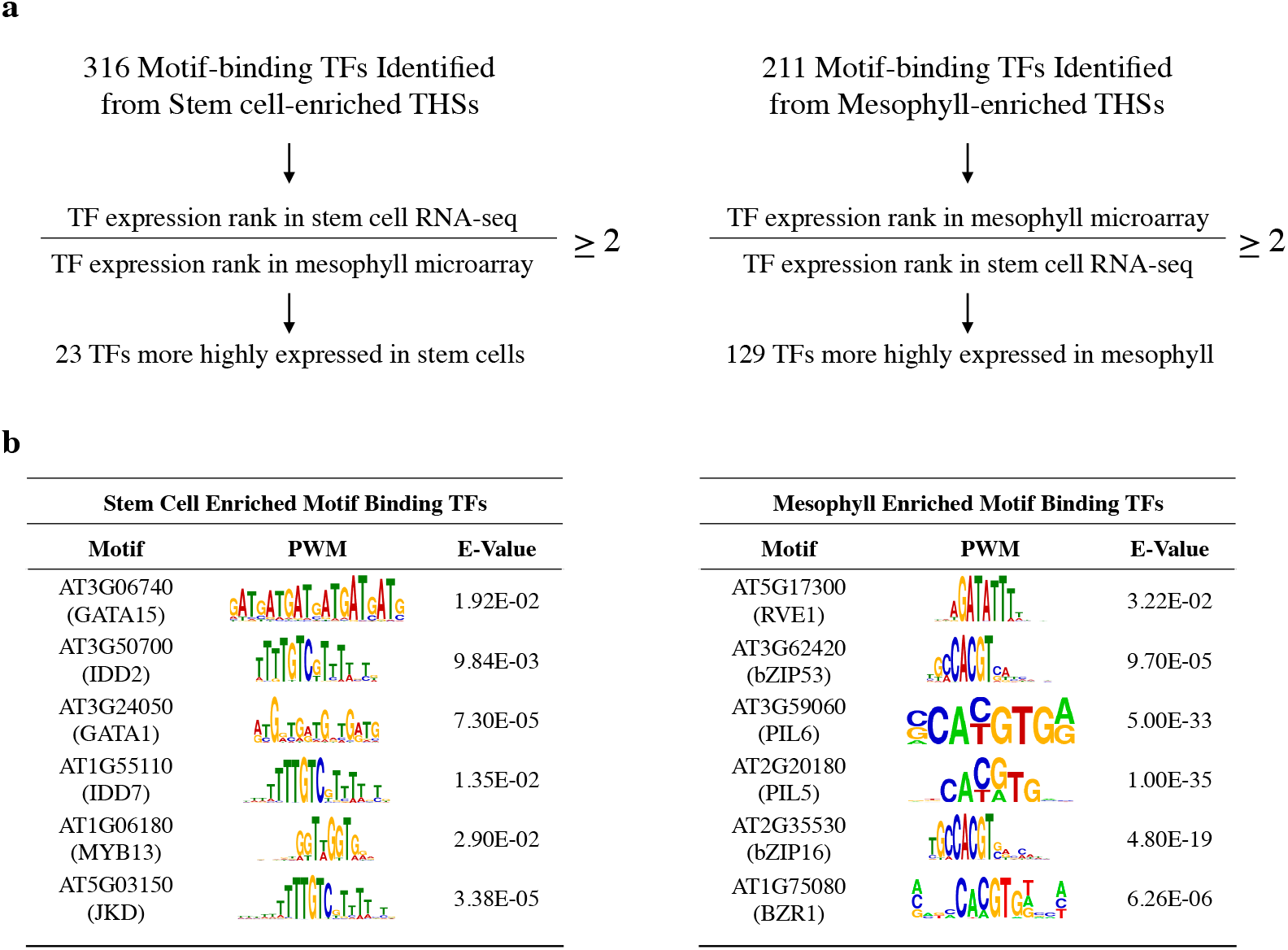
Sequence motifs identified in cell type-enriched THSs. **(a)** Cell type-enriched THS sequences were centered and scaled to 300 bp, repeat masked, and analyzed with MEME-ChIP (Methods). Motifs that had an E-value equal to or less than 0.05 were considered significant. The 316 and 211 transcription factors (TFs) associated with overrepresented motifs from stem cell-and mesophyll-enriched THSs, respectively, were further separated by their ranked expression difference between previously reported stem cell RNA-seq and mesophyll microarray data [28–29]. Only those TFs that had at least a two-fold higher expression difference for the cell type their motif was identified in were kept (Table S5). **(b)** Six TFs that potentially regulate transcriptional networks for each cell type, their position weight matrix, and E-value from the MEME-ChIP analysis are shown for the stem cell (left) and mesophyll (right). The TFs are ranked by their difference in expression between the two cell types, with the highest expression difference for the corresponding cell type at the top.

Using this approach, we discovered a putative stem cell-specific functional network of 5 interconnected TFs that belong to two distinct families: INDETERMINATE DOMAIN C2H2 zinc finger protein family (IDD) and GATA factor zinc finger transcription factor protein family (Figure 3B and Figure S4A). The *Arabidopsis* IDD family of TFs has 16 members, which are involved in promoting gibberellin signaling, auxin biosynthesis and transport, and lateral organ differentiation, but are best known for their control of tissue formation during root development [32–34]. The GATA TF family is comprised of approximately 30 members, which can be divided into four subfamilies [35]. Of these subfamilies, the best studied TFs are the members of the B-GATA subfamily, including GNC and its paralog GNL, which are involved in the control of greening and regulation of plant development downstream of the hormones gibberellin, cytokinin, and auxin [36–38].

We carried out FIMO analysis [39] using all the stem cell THS sequences to identify motif occurrences and thus predicted binding sites for each of the four TFs from the STRING-derived regulatory network: INDETERMINATE DOMAIN 2 (IDD2), INDETERMINATE DOMAIN 7 (IDD7), GATA TRANSCRIPTION FACTOR 1 (GATA1), and GATA TRANSCRIPTION FACTOR 15 (GATA15). These predicted binding sites were then used to locate the nearest TSS to identify the set of predicted target genes for each of the four TFs (Figure 4A and B). Using this approach, we discovered 3,218 predicted target genes for GATA15, 5,946 for IDD2, 3,603 for GATA1, and 5,322 for IDD7. Out of the 9,962 target genes for these TFs, 569 genes are predicted targets for all four TFs. We performed GO analysis on this group of genes using AgriGO (Figure 4C). The GO terms overrepresented in this analysis revealed that many of the target genes predicted to be regulated by IDD and GATA TFs are involved in control of auxin-mediated signaling, regulation of transcription, and shoot development (Figure 4C). A STRING network of interactions among these target genes, under high stringency (a minimum interaction score of 0.700), is shown in Figure S5. Notably, we found that the known stem cell regulator CLV3 is a target of this IDD/GATA gene regulatory network.

**Figure 4.**
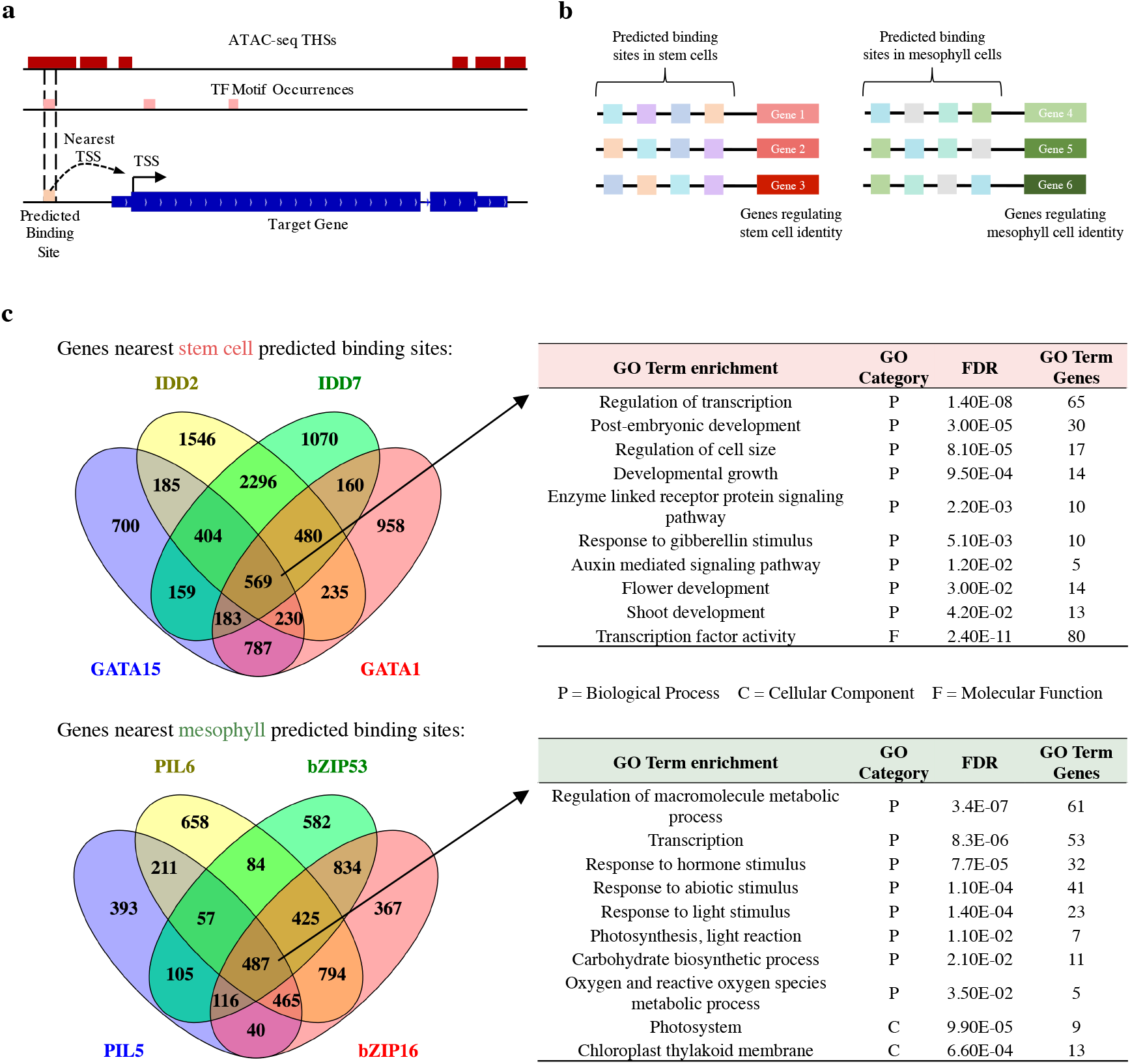
Predicted binding sites and target genes for cell type-enriched TFs. **(a)** Schematic for identifying predicted binding sites using ATAC-seq THSs and HMO-identified TF motif occurrences in the genome. These predicted sites were used to identify the nearest TSS to define the target gene potentially regulated by the TF. (b) Schematic for using the predicted binding sites (as shown in **a**) to identify genes that regulate cell identity, and which TFs control the expression of these genes. (c) Overlap of predicted target genes of IDD2, IDD7, GATA15, and GATA1 (top, left). The genes targeted by all four TFs (569) were analyzed with AgriGO, and the resulting GO terms that had an FDR value of 0.05 or less were retained. A subset of these enriched GO terms are shown (top, right). Overlap of predicted target genes of PIL6, PIL5, bZIP53, and bZIP16 (bottom, left). The genes targeted by all four factors (487) were analyzed with AgriGO, and the resulting GO terms that had an FDR value of 0.05 or less were retained. A subset of these enriched GO terms are shown (bottom, right).

The 129 mesophyll-enriched TFs whose motifs were overrepresented in the mesophyll-enriched THSs (Figure 3A) had a high density of functional interconnections when analyzed with the STRING database (Figure S4B). We identified three major mesophyll-specific sub networks of TFs. The largest sub network was comprised of 41 extensively interconnected TFs, including 10 members of WRKY and 11 members of ERF family of TFs, which are known to regulate various biotic and abiotic stress responses [40–46]. Seven out of the eight TFs in the second sub network belong to the TEOSINTE BRANCHED 1, CYCLOIDEA, PCF1 (TCP) family, which is known to control plant growth and organ development, including leaf development [47–50]. The third sub network included eight well-connected TFs. Among these are three members of the PIF family: PHYTOCHROME INTERACTING FACTOR 3-LIKE 5 (PIL5), PHYTOCHROME INTERACTING FACTOR 3-LIKE 6 (PIL6), and PHYTOCHROME-INTERACTING FACTOR 7 (PIF7). Two additional TFs found in this subnetwork, BES1-INTERACTING MYC-LIKE 1 (BIM1) and BRASSINAZOLE-RESISTANT 1 (BZR1), are involved in the brassinosteroid (BR) hormone signaling pathway. PIFs belong to the bHLH family of TFs and are best known as negative regulators of chlorophyll biosynthesis and photomorphogenesis [51–54]. BRs are important regulators of many aspects of plant growth and developmental processes including cell elongation, responses to biotic and abiotic stresses, and photomorphogenesis [55–56]. We decided to explore this PIF/BR regulatory sub network in more detail since both PIFs and BRs have been implicated in the regulation of chloroplast biogenesis [51–54–57], and therefore may directly affect the physiology and development of mesophyll cells.

As with the IDD/GATA regulatory network in the stem cells, our next goal was to identify the putative target genes of TFs enriched in mesophyll. In this case, we also included two additional TFs, out of the 129 mesophyll-enriched TFs, that belong to the bZIP family: bZIP16, and bZIP53. We chose to include bZIP TFs because it has been recently shown that PIF and bZIP TFs HY5 and HYH interact with each other and form heterodimers to antagonistically regulate chlorophyll biosynthesis [58–59] and the production of Reactive Oxygen Species (ROS) during deetiolation [58].

Using mesophyll-enriched THS sequences we performed FIMO analysis [39] to identify predicted binding sites for each of four TFs of interest: PIL5, PIL6, bZIP16, and bZIP53. We then identified putative target genes by assigning each predicted binding site to its nearest TSS. As seen for the stem cell TFs, all four of the mesophyll-enriched TFs also showed extensive co-regulation of common target genes. We then performed GO analysis on the set of 487 target genes putatively regulated by all four TFs (Figure 4B). Many of the GO terms identified describe known functions of mesophyll cells including response to abiotic stimulus and light stimulus, photosynthesis-light reaction, and carbohydrate biosynthetic process (Figure 4C and Table S6). These results suggest that the PIL5, PIL6, bZIP16, and bZIP53 TFs likely play important roles in regulating mesophyll physiological functions.

We discovered that many of the putative target genes of the IDD and GATA TFs in stem cells, and PIFs and bZIPs in mesophyll cells, are TFs themselves. This finding alludes to the presence of cell type-specific transcription factor cascades, which may regulate important biological processes in these two cell types (GO term: regulation of transcription, Figure 4C). To explore TF-to-TF connections in greater detail, we explored putative regulatory connections among the stem cell and mesophyll TFs to illuminate how they might regulate each other. Previous studies have used similar models with great success in order to build de-novo TF regulatory networks for 41 human cell types [60]. The model presented in Figure 5A describes the logic of this analysis, in which each TF can bind to its recognition motif found within its own regulatory regions and/or within the regulatory regions of other TF genes. For instance, the proximal regulatory region of a hypothetical transcription factor gene (TF5) gene contains DNA-binding motifs of four other TFs: TF1, TF2, TF3, and TF4. The DNA-binding motif of the TF5 is, on the other hand, found in the upstream region of TF4, which also has its recognition motif present in the regulatory regions of TF1 and TF2 (Figure 5A). Thus, an extensive co-regulatory network of multiple TFs can be mapped in this manner.

**Figure 5.**
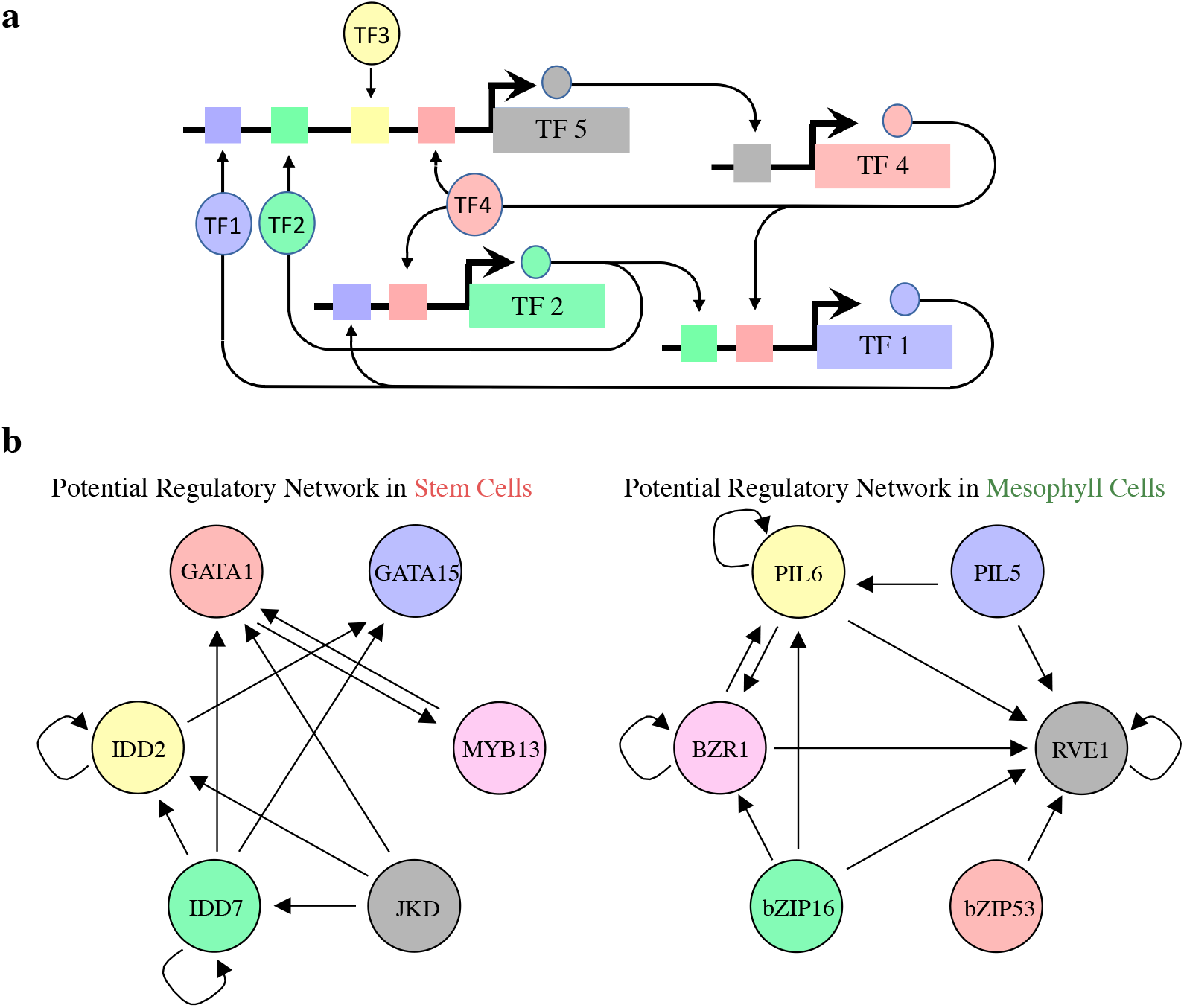
Proposed regulatory pathways for key transcription factors in stem cells and mesophyll cells. **(a)** Schematic for identifying regulatory interactions between transcription factors (TFs). A predicted binding site for a TF, such as TF5, may regulate the expression of another TF, such as TF4. Subsequently regulated TFs may regulate other TFs, making up a transcription factor network that is active within a cell type. **(b)** The putative regulatory networks for stem cells (left) and mesophyll (right) are shown. Each TF circle has regulatory inputs (stem cell or mesophyll predicted TF binding site within its proximal regulatory regions) and regulatory outputs (that TF’s predicted binding site in the other TF gene’s proximal regulatory regions). For example, IDD7 has four regulatory outputs to IDD2, GATA1, GATA15, and itself, and one regulatory input to itself.

Using the strategy described in Figure 5A for the predicted target genes for each TF, we derived more comprehensive stem cell-specific and mesophyll-specific putative regulatory circuitries of TFs, further uncovering complex combinatorial interactions among TFs within these networks (Figure 5B). For example, in the stem cell-specific TF network, IDD7 appears to regulate itself and three other TFs: IDD2, GATA1, and GATA15, but not JKD or MYB13 (Figure 5B). On the other hand, JKD may regulate the expression of IDD2, IDD7, and GATA1, but not that of GATA15 and MYB13. GATA1 and MYB13 seem to regulate each other, while GATA15 appears to be most downstream component in this TF hierarchy since it does not regulate any other TF in this network.

Similarly, in the mesophyll-specific TF regulatory network, PIL6 and BZR1 seem to regulate both themselves and each other. bZIP16 appears to be at the apex of this regulatory module since it regulates three different TFs, while all others regulate the expression of two or fewer different TFs. Importantly, this model predicts that PIL6 and bZIP16, as well as PIL5 and bZIP53, co-regulate the expression of RVE1, which resembles the coordinated TF interaction previously described for another pair of bHLH/bZIP TFs: PIFs and HY5/HYH [58–59].

## Conclusions

### Cell type-specific THSs contain cis-regulatory elements relevant to the physiology of stem cells and mesophyll cells

Since fully differentiated mesophyll cells in leaves are at the very end of the differentiation process from stem cells in the meristem, it was perhaps surprising to find that more than 91% of the reproducible mesophyll THSs identified by peak calling alone were also present in stem cells (Figure 1C). These results indicated that the accessible chromatin regions of these two cell types are not as different as we originally anticipated. Nevertheless, we were able to identify several thousand cell type-unique THSs (Figure 1C) that were only detected in one cell type or the other. Since the majority of THSs were shared between the two cell types, we performed a quantitative analysis to identify THSs that were differentially accessible between stem cells and mesophyll cells. This analysis led to the identification of several thousand more THSs in each cell type than were identified by the absolute, all-or-nothing, peak calling strategy. We assigned each of these cell type-enriched THSs to their nearest TSS as the putative target gene regulated by the differential accessibility event. Gene Ontology (GO) analysis of the stem cell THS-proximal genes identified overrepresented GO terms that describe known functions of the SAM stem cells in regulating cell differentiation and organ development (Figure 2D). Similarly, we identified GO terms for the mesophyll THS-proximal genes that are consistent with established roles of mesophyll cells in mediating the responses of various biotic and abiotic stresses (Figure 2D).

Overall, these results indicate that INTACT-ATAC-seq allows us not only to successfully identify cell type-enriched THSs, but also that these THSs likely contain regulatory elements that are highly relevant for the biology of these two cell types.

### Mesophyll-enriched THSs are already accessible in stem cells

In comparing open chromatin regions between cell types, we discovered that the stem cell-enriched THSs tend to be much more highly accessible in stem cells relative to mesophyll, but that these regions still showed a low level of accessibility in the mesophyll cell type (Figure 3B). This is consistent with our previous observation that the root epidermal hair and non-hair cell types show mainly quantitative, rather than qualitative differences in chromatin accessibility [23]. These results also suggest that, at least in the *Arabidopsis* cell types examined, a given regulatory region is never completely inaccessible in any cell type, and this may simply reflect the proportion of cells in the population in which a TF binding event is occurring at that location.

When we examined chromatin accessibility at mesophyll-enriched THSs, we found that while accessibility was far higher in mesophyll cells, the stem cells also showed significant accessibility at these sites (Figure 3B). Thus, while stem cell-enriched THSs represent regions that are highly accessible in stem cell and far less so in mesophyll, the mesophyll-enriched THSs tend to already be highly accessible in the progenitor stem cells. This suggests that even mesophyll cell-enriched THSs are available for TF binding in stem cells, and this phenomenon could underlie the developmental flexibility of stem cells. Whether this observation is a unique characteristic of the SAM stem cell chromatin or more universal feature of any plant stem cell chromatin in comparison to differentiated cells remains to be tested. Regardless, we can hypothesize that one of the reasons why pluripotent stem cells in the SAM would maintain more accessible regulatory elements is to allow them flexibility to change their transcriptome in response to stimuli. In other words, by being more open, the stem cell chromatin is more “primed” for transcriptional reprogramming, thereby endowing stem cells with the plasticity to efficiently respond to differentiation cues.

### Identified cell type-specific transcriptional modules are likely important for the establishment of lineage-specific regulatory programs in each cell type

In a search for TFs that should be relevant to the biology of each cell type, we analyzed the differentially enriched THS regions to identify putative cell type-specific *cis-*regulatory motifs as well as the TFs that bind them (Figure 3A). Using publicly available expression data for these two cell types, we found TFs that were differentially expressed in each cell type and whose motifs were also overrepresented in THSs enriched in that cell type. We analyzed these cell type-enriched TFs using the STRING database to identify modules of TFs that might act coordinately in each cell type (Figure 3B).

Following the logic that the identified TF motifs likely represent true TF binding events when they occur within an open chromatin region of the corresponding cell type, we were able to predict the target genes of TFs of interest (Figure 4A). We found that in each cell type, cell type-enriched TFs showing connections in the STRING database also tended to co-regulate many genes (Figure 4C). In each case, a relatively large gene set appeared to be co-regulated by all four TFs, and GO analysis of these gene sets revealed functions consistent with the biology of each cell type. Thus, while using predicted target genes, rather than direct measurement of TF binding by ChIP-seq, may lead to the inclusion of false positive binding events, there is strong evidence that many true positives exist among the putative target genes.

The TF modules we identified in each cell type by expression and STRING analysis were then used to define regulatory interactions between cell type-specific TFs (Figure 5B). The predicted combinatorial interactions among TFs within these regulatory networks were extensive and likely play important roles in establishing and/or maintaining cell type-specific transcriptional programs during differentiation. These new hypotheses can now be experimentally tested. For instance, the STRING-derived stem cell TF network was comprised of the members of IDD and GATA TF families. While the functions of individual members of IDD and GATA families of TFs are more or less known, to our knowledge, the functional interactions between GATA and IDD TFs have never been proposed or studied. In addition, IDDs are known to regulate lineage identity, patterning, and formative divisions throughout *Arabidopsis* root growth [34] but have not been implicated in a similar role during vegetative meristem development and differentiation, which our data now suggest. Interestingly, *CLV3* itself was identified as a target gene of the IDD/GATA regulatory network (Figure S5), further supporting our hypothesis that this regulatory circuitry may play an important role in stem cell homeostasis. One way to test these hypotheses is by manipulating the expression of the IDD/GATA TFs specifically in the SAM stem cells. For instance, the *CLV3* regulatory sequences can be used to drive the expression of RNAi or artificial microRNA constructs to specifically knock down the IDD/GATA TFs in the stem cell population. In addition, an inducible overexpression system may be utilized to overproduce IDD/GATA TFs specifically in the SAM in order to monitor chromatin and transcriptional changes. The results from these experiments will address whether the IDD/GATA TFs indeed act as important regulators of SAM function, and further characterize these regulatory connections.

It has been previously demonstrated that the PIF and bZIP transcription factors HY5 and HYH antagonistically regulate chlorophyll biosynthesis during seedling development [58–59]. Our results now indicate that other members of the PIF and bZIP families of TFs may cooperate in regulating chlorophyll biosynthesis and light responses in mesophyll cells. This intricate cooperation between two TFs with potentially opposite functions may allow mesophyll cells to fine-tune their transcriptional programs in response to various hormonal and environmental stimuli during leaf development and/or throughout daily light/dark cycles. Experimental manipulations to modulate the expression of mesophyll PIFs and bZIPs by overexpressing or suppressing these TFs specifically in mesophyll cells can now be performed to test these new hypotheses.

### INTACT-ATAC-seq as a powerful technique for predicting cell type-specific transcriptional regulatory networks

In this study, we combined the INTACT method with ATAC-seq to successfully isolate nuclei from two specific cell types and locate differentially accessible chromatin regions containing important cis-regulatory motifs. This allowed us to identify the TFs that likely bind at these regulatory elements and to construct cell type-specific TF regulatory networks. Our data provide new hypotheses and will serve as a valuable resource that can be used to derive further de-novo models of transcriptional regulatory networks relevant to cell fate specification during differentiation. These hypotheses can be experimentally tested and the results from these experiments used to further build upon and expand our current understanding of the regulatory mechanisms controlling cell fate and function during plant development.

## Methods

### Plant growth conditions and transformation

*Arabidopsis thaliana* plants of the Columbia (Col-0) ecotype were grown in soil or on half-strength Murashige and Skoog (MS) media [61] agar plates in growth chambers under fluorescent lights, with 16 hour light-8 hour dark cycle at 20°C. All seeds, either on agar plates or in the soil, were stratified for three days at 4°C prior to moving them to the growth chambers. Plasmid constructs were introduced into *Agrobacterium tumefaciens GV3101* strain by electroporation. Plant transformation was performed using the floral dip method with corresponding *Agrobacterium* clones [62]. Primary transformant seedlings (*T*_1_) were first selected on half-strength MS media agar plates containing 35mg/L hygromycin, 25mg/L glufosinate ammonium (BASTA), and 100mg/L timentin, and then transferred to soil.

### Plasmid DNA constructs

We used the promoters of *CLAVATA3 (CLV3)* and *Rubisco small subunit 2B (RBC)* genes, known to be exclusively transcribed in stem cells and mesophyll cells, respectively [17–19], to drive cell-type specific expression of the *NTF* gene. To construct the *CLV3p::NTF* plasmid, the *NTF* coding sequence [15] was first PCR amplified using the forward primer 5′-cat<u>ctgcag</u>atgaatcattcagcgaaaacc-3′, introducing a *PstI* restriction site (underlined), and the reverse primer 5′-cat<u>ggatcc</u>tcaagatccaccagtatcctc-3′, introducing a *BamHI* restriction site (underlined). The PCR product was then digested with *PstI/BamHI* enzymes and ligated into *PstI/BamHI* sites of the *pBU14* plasmid containing the *CLV3* promoter and terminator sequences [63]. The *ACT2p::BirA* plasmid has been previously described [64]. The *RBCp::NTF* construct was produced by removing the *ADF8* promoter from the previously described *ADF8p::NTF* plasmid [15] via *XmaI* and *NheI,* and replacing it with a 1.5 kb upstream fragment of *RBC*, including the start codon.

### Microscopy

The shoot apical meristems of six days old *T*_3_ *CLV3p::NTF;ACT2p::BirA* seedlings and leaves 5 and 6 of three weeks old *T*_3_ *RBCp::NTF;ACT2p::BirA* plants were observed using a Leica SP8 confocal laser scanning microscope with 42x immersion objectives to confirm proper expression and localization of the NTF protein. GFP fluorescence was visualized by excitation at 488 nm. The shoot apical meristems were visualized by manually removing the surrounding leaf tissue and imaging the shoot apical meristem from the side. Mesophyll cells were visualized by dissecting the leaf with a scalpel and imaging the cross section. For each sample, the tissue was immersed in perfluoroperhydrophenanthrene, covered with a cover slip, and imaged.

### Nuclei isolation by INTACT

Purification of nuclei from specific cell types using the Isolation of Nuclei TAgged in specific Cell Types (INTACT) method was performed as described previously [20] with following modifications: 0.5 grams of freshly harvested plant tissue was used for nuclei isolation; *CLV3p::NTF;ACT2p::BirA* transgenic seedlings were collected at 6 days of age and were processed by grinding the tissue to a fine powder using liquid nitrogen. Leaves 5 and 6 from three week old plants with the *RBCp::NTF;ACT2p::BirA* transgenics were collected and finely chopped with a razor blade in Nuclei Purification Buffer (NPB) on ice. For both preparations, a volume of 10 εl of Streptavidin M280 magnetic beads was used to capture biotinylated nuclei.

Compared to previous INTACT-ATAC-seq experiments using root tissue, we observed a much higher level of organelle contamination in purified nuclei from shoot tissue of both the *CLV3p::NTF;ACT2p::BirA* and *RBCp::NTF;ACT2p::BirA* transgenic lines, as revealed by the large percentage of organelle-derived reads in our datasets. During mesophyll nuclei purification in particular, we observed many large clusters of nuclei associated with magnetic beads, suggesting that chloroplasts and mitochondria may become trapped within these clusters. Further optimization of the INTACT procedure on green tissue will likely eliminate this issue. For instance, it may be necessary to use even smaller amounts of starting tissue, to further decrease the amount of streptavidin beads used in order to decrease bead clustering, and to add additional washing steps or use higher non-ionic detergent concentrations during purification.

### Assay for transposase accessible chromatin (ATAC) and library preparation

It is important to note that all INTACT-purified nuclei were isolated and used fresh, and were never frozen prior to the transposase integration reaction. Transposase tagmentation and sequencing library preparations were then carried out as previously described [20]. Briefly, 25,000 purified nuclei were resuspended in a 50 μl transposase integration reaction and incubated at 37° C for 30 min using Nextera reagents (Illumina, FC-121-1030). Tagmented DNA was purified using the MiniElute PCR purification kit (Qiagen), eluted in 11 μl of elution buffer, and then the sample was amplified using 2X high fidelity PCR mix (NEB) with custom barcoded primers for 10–12 total PCR cycles. The amplified ATAC-seq libraries were purified using AMPure XP beads (Beckman Coulter) and then quantified by qPCR using the NEBNext library quant kit (NEB). The quantified libraries were analyzed using a Bioanalyzer high sensitivity DNA chip (Agilent) before pooling and next-generation sequencing.

### High throughput sequencing

Next-generation sequencing was done using the NextSeq 500 instrument (Illumina) at the Georgia Genomics Facility at the University of Georgia. All libraries were pooled and sequenced in the same flow cell using paired-end 36 nt reads.

### Sequence read mapping, processing, and visualization

Sequencing reads were mapped to the *Arabidopsis thaliana* genome (version TAIR10) using Bowtie2 software [65] with default parameters. Mapped reads in *.sam* format were converted to *.bam* format and sorted using Samtools 0.1.19 [66]. Mapped reads were filtered using Samtools to retain only those reads that had a mapping quality score of 2 or higher (Samtools *“view”* command with option “-q 2” to set mapping quality cutoff). These reads were further filtered with Samtools to keep only the reads that mapped to nuclear chromosomes, thereby removing reads that mapped to either the chloroplast or mitochondrial genomes. Finally, the stem cell and mesophyll cell datasets were also processed such that the experiments within a biological replicate had the same number of mapped reads prior to further analysis (Samtools “*view*” command with option “-*c*” to count the number of aligned reads in each dataset and “-*S*” to scale down by the numerical fraction the number of aligned reads to be kept). For visualization, the filtered, sorted, and scaled .*bam* files were converted to the bigwig format using the *“bamcoverage”* script in deepTools 2.0 [21] with a bin size of 1 bp and RPKM normalization. Heatmaps and average plots displaying ATAC-seq data were also generated using the “*computeMatrix*” “*plotHeatmap*” and “*plotProfile*” functions in the deepTools package. Genome browser images were made using the Integrative Genomics Viewer (IGV) 2.3.68 [67] with bigwig files processed as described above.

### Peak calling to detect transposase hypersensitive sites (THSs)

Peak calling on ATAC-seq data was performed using the “*Findpeaks*” function of the HOMER package [22] with the parameters “-*minDist 150*” and “-*region*”. These parameters set a minimum distance of 150 bp between peaks before they are merged into a single peak and to allow identification of variable length peaks, respectively. We refer to the peaks called in this way as “transposase hypersensitive sites,” or THSs. To deepen our analysis and increase the resolution and number of THSs called in the two cell types we utilized an additional parameter when comparing the degree of accessibility between the two cell types. The additional parameter “ -*regionRes* 1” separated larger THSs into several smaller THSs without affecting the way in which THSs that were several hundred base pairs in size or smaller are called. For calling peaks in genomic DNA we similarly employed the “*Findpeaks*” function using the parameters “-*minDist 150*” and “-*region*”.

### Genomic distribution of THSs

The distribution of THSs relative to genomic features was identified using the PAVIS web tool [68] with “upstream” regions set as the 2,000 bp upstream of the annotated transcription start site, and “downstream” regions set as the 1,000 bp downstream of the transcript end site.

### THSs enriched in a specific cell type

The number of reads (counts) present in each cell type at all the THSs called in stem cell and mesophyll ATAC-seq data was obtained using HTSeq’s *htseq-count* script [69]. Two replicates of each cell type were counted and the counts were processed using DESeq2 [24]. THSs that had an adjusted p-value ≤ 0.05 and log fold change of 1 or more for a specific cell type were identified as THSs enriched in that cell type.

### Transcription factor motif analysis

ATAC-seq THSs that were enriched in one cell type or the other were used for motif analysis. The cell type-enriched THSs from each cell type were first adjusted to the same size (300 bp). The sequences present in these scaled regions were isolated using the Regulatory Sequence Analysis Tools (RSAT), which also masks any repeat sequences [70]. The masked sequences were run through MEME-ChIP with default parameters to identify motifs that were present in higher proportions than expected by chance (i.e. overrepresented motifs) [28]. The DREME, MEME, and CentriMo programs were used to identify overrepresented motifs, and Tomtom matched these motifs to previously reported TF binding motifs. Motifs from both Cis-BP [71] and DAP-seq [72] databases were used in all motif searches, and only those that had an E-value < 0.05 were considered significant.

### Assignment of ≤ to nearby genes

For each ATAC-seq dataset, the THSs were assigned to putative target genes using the “TSS” function of the PeakAnnotator 1.4 program [25]. This program assigns each THS to the closest transcription start site regardless of whether it is upstream or downstream from the THS, and reports the distance from the peak center to the TSS based on the genome annotations described above.

### Publicly available RNA-seq and microarray data

Published RNA-seq data from the CLV3-expressing cell population of shoot meristems [29], isolated from 21 day old plants, and microarray data from mesophyll cells isolated at ZT04 from 10 days old cotyledons grown under a long day (LD) cycle (GSM1219271, [30]), were used to define TFs that were differentially expressed in the stem cells relative to the mesophyll cells, and vice versa.

### Calculating the relative expression ranks of TFs in RNA-seq and microarray data sets

Within the stem cell RNA-seq dataset, the genes were considered expressed if the FPKM value was ≥1, and 17,811 genes satisfied this criterion. Within the mesophyll microarray data, there were 28,583 expressed genes. For each data set, we first arranged the genes based on the level of their expression from highest to lowest. Next, for each TF of interest we calculated the percentile of its expression relative to total number of expressed genes in each data set and then measured the difference in its expression rank between the two cell types (Table S5). For instance, GATA15 TF ranked as the 2085th most highly expressed gene in the stem cell RNA-seq data set, which equals 11.7% (2,085/17,811) in expression ranking. The same TF in the mesophyll expression data set ranked 24,755^th^ most highly expressed, which is 86% (24,755/28,583) in expression ranking. We then calculated the difference in expression ranking of GATA15 between the two cell types by dividing the relative expression ranks in percentages (11.7/86). TFs that have at least a two fold difference in their relative expression ranking between cell types were considered as more highly expressed in one cell type or the other.

### Protein interaction analysis using STRING

Gene lists were analyzed using the STRING database to identify groups of TFs that have predicted interactions based on the co-expression analysis, publication co-occurrences, colocalization, gene orthology, and experimental information such as yeast-2-hybrid interactions [31]. The network connections between the submitted TFs were visualized by their confidence score, where a thicker line indicates a higher interaction score. Furthermore, the network was subdivided into differentially colored nodes by the Markov Cluster Algorithm score set to 3.0. This allows for the detection of genes with some evidence for interactions, but whose association does not pass the interaction threshold required to have a bona fide connection. The scale of interaction scores in STRING is as follows: 0.15=low confidence, 0.4=medium confidence, 0.7=high confidence, and 0.9=highest confidence. The minimum interaction threshold used in this study was set to at least 0.400 or 0.700. The inputs used for the STRING database were the *Arabidopsis* gene IDs.

### Defining predicted binding sites for transcription factors

We used FIMO [39] to identify TF motif occurrences within the repeat-masked sequence of the *Arabidopsis* genome. Significant motif occurrences were those with a p-value < 0.0001. Predicted binding sites for a given TF were defined as motif occurrences that were present within THSs of a given cell type (see Figure 4A for a schematic diagram of this process).

### Gene ontology analysis

Gene ontology (GO) analysis was carried out on gene lists using the AgriGO GO Analysis Toolkit, with default parameters [26–27]. GO terms that had a false discovery rate (FDR) of 0.05 or less were considered significant.

## Additional files

List of Supplemental figures and tables and a short description of each

Additional file 1: Figure S1. ATAC-seq read alignments, sample variability, and fragment size distribution.

Additional file 2: Table S1. THS coordinates in cell type replicates, genomic DNA, enriched THSs, and higher resolution peak calling.

Additional file 3: Table S2. DESeq2 results for counts obtained in two replicates of each cell type.

Additional file 4: Figure S2. Genomic distribution of cell type-enriched THSs and chromatin profiles of nearby genes.

Additional file 5: Table S3. AgriGO results for genes nearest to cell type-enriched THSs.

Additional file 6: Figure S3. Identification of THSs in genomic DNA.

Additional file 7: Table S4. MEME-ChIP results for cell type-enriched THS sequences.

Additional file 8: Table S5. Transcription Factor expression differences analyzed by comparing expression rank change between previously published stem cell RNA-seq and mesophyll microarray data.

Additional file 9: Figure S4. Interactions among TFs enriched in each cell type.

Additional file 10: Table S6. Coordinates of predicted binding sites in the two cell types, the nearest genes they likely regulate, and AgriGO results for genes with predicted binding sites for all four TFs, for each cell type.

Additional file 11: Figure S5. Predicted regulatory networks for stem cell and mesophyll cells.

## List of Abbreviations

(SAM): Shoot apical meristem
(INTACT): Isolation of Nuclei TAgged in specific Cell Types
(ATAC-seq): Assay for Transposase-Accessible Chromatin with high-throughput sequencing
(THS): Transposase Hypersensitive Site
(TF): Transcription Factor

## Declarations

### Availability of Data and Material

The raw and processed ATAC-seq data described in this work has been deposited to the NCBI Gene Expression Omnibus (GEO) database under the record number GSE101940. All transgenic lines used in this study are freely available by request.

### Competing Interests

The authors have no competing interests to declare.

### Funding

This work was made possible by start-up funding from Emory University and a grant from the National Science Foundation (Plant Genome Research Program grant #IOS-123843).

### Authors’ Contributions

MB, PS, and RBD designed the research project. PS produced the stem cell INTACT line, purified all nuclei, and made the ATAC-seq libraries. MB performed the majority of data analyses, with assistance from RBD. PS contributed to identifying TFs enriched in one cell type or the other. E.C.M and R.B.M. generated the mesophyll INTACT construct, produced a transgenic line homozygous for both *RBCp:NTF and ACT2p:BirA* transgenes, and performed the initial characterization of the line. MB and PS drafted the manuscript with editing from R.B.D.

#### Acknowledgements

We would like to thank Kelsey Maher for constructive criticism of the manuscript. Funding for this work was provided by the National Science Foundation and Emory University.

### Ethics Approval and Consent to Participate

Not Applicable

### Consent for Publication

Not applicable

## References

1 Besnard F, Vernoux T, Hamant O. Organogenesis from stem cells in planta: multiple feedback loops integrating molecular and mechanical signals. Cell Mol Life Sci. 2011;68:2885–906.

2 Barton MK. Twenty years on: the inner workings of the shoot apical meristem, a developmental dynamo. Dev Biol. 2010;341:95–113.

3 Spitz F, Furlong EE. Transcription factors: from enhancer binding to developmental control. Nat Rev Genet. 2012;13:613–26.

4 Burton A, Torres-Padilla ME. Chromatin dynamics in the regulation of cell fate allocation during early embryogenesis. Nat Rev Mol Cell Biol. 2014;15:723–34.

5 Schiefelbein J, Huang L, Zheng X. Regulation of epidermal cell fate in Arabidopsis roots: the importance of multiple feedback loops. Front Plant Sci. 2014;5:47.

6 Balcerowicz D, Schoenaers S, Vissenberg K. Cell Fate Determination and the Switch from Diffuse Growth to Planar Polarity in Arabidopsis Root Epidermal Cells. Front Plant Sci. 2015;6:1163.

7 John S, Sabo PJ, Canfield TK, Lee K, Vong S, Weaver M, et al. Genome-scale mapping of DNase I hypersensitivity. Curr Protoc Mol Biol. 2013;Chapter 27: Unit 21.7.

8 He HH, Meyer CA, Hu SS, Chen MW, Zang C, Liu Y, et al. Refined DNase-seq protocol and data analysis reveals intrinsic bias in transcription factor footprint identification. Nat Methods. 2014;11:73–8.

9 Zhong J, Luo K, Winter PS, Crawford GE, Iversen ES, Hartemink AJ. Mapping nucleosome positions using DNase-seq. Genome Res. 2016;26:351–64.

10 Sheffield NC, Thurman RE, Song L, Safi A, Stamatoyannopoulos JA, Lenhard B, et al. Patterns of regulatory activity across diverse human cell types predict tissue identity, transcription factor binding, and long-range interactions. Genome Res. 2013;23:777–88.

11 Thurman RE, Rynes E, Humbert R, Vierstra J, Maurano MT, Haugen E, et al. The accessible chromatin landscape of the human genome. Nature. 2012;489:75–82.

12 Buenrostro JD, Giresi PG, Zaba LC, Chang HY, Greenleaf WJ. Transposition of native chromatin for fast and sensitive epigenomic profiling of open chromatin, DNA-binding proteins and nucleosome position. Nat Methods. 2013;10:1213–8.

13 Buenrostro JD, Wu B, Chang HY, Greenleaf WJ. ATAC-seq: A Method for Assaying Chromatin Accessibility Genome-Wide. Curr Protoc Mol Biol. 2015;109:21.9.1-9.

14 Lu Z, Hofmeister BT, Vollmers C, DuBois RM, Schmitz RJ. Combining ATAC-seq with nuclei sorting for discovery of cis-regulatory regions in plant genomes. Nucleic Acids Res. 2016.

15 Deal RB, Henikoff S. A simple method for gene expression and chromatin profiling of individual cell types within a tissue. Dev Cell. 2010;18:1030–40.

16 Deal RB, Henikoff S. The INTACT method for cell type-specific gene expression and chromatin profiling in Arabidopsis thaliana. Nat Protoc. 2011;6:56–68.

17 Schoof H, Lenhard M, Haecker A, Mayer KF, Jurgens G, Laux T. The stem cell population of Arabidopsis shoot meristems in maintained by a regulatory loop between the CLAVATA and WUSCHEL genes. Cell. 2000;100:635–44.

18 Yadav RK, Girke T, Pasala S, Xie M, Reddy GV. Gene expression map of the Arabidopsis shoot apical meristem stem cell niche. Proc Natl Acad Sci U S A. 2009;106:4941–6.

19 Sawchuk MG, Donner TJ, Head P, Scarpella E. Unique and overlapping expression patterns among members of photosynthesis-associated nuclear gene families in Arabidopsis. Plant Physiol. 2008;148:1908–24.

20 Bajic M, Maher KA, Deal RB. Identification of Open Chromatin Regions in Plant Genomes Using ATAC-Seq. Methods Mol Biol. 2018;1675:183–201.

21 Ramirez F, Ryan DP, Gruning B, Bhardwaj V, Kilpert F, Richter AS, et al. deepTools2: a next generation web server for deep-sequencing data analysis. Nucleic Acids Res. 2016;44:W160–5.

22 Heinz S, Benner C, Spann N, Bertolino E, Lin Y C, Laslo P, et al. Simple combinations of lineage-determining transcription factors prime cis-regulatory elements required for macrophage and B cell identities. Mol Cell. 2010;38:576–89.

23 Maher KA BM, Kajala K, Reynoso M, Pauluzzi G, West DA, et al. bioRxiv.org – the preprint server for Biology. https://www.biorxiv.org/content/early/2017/07/24/167932. Accessed 24 July 2017.

24 Love MI, Huber W, Anders S. Moderated estimation of fold change and dispersion for RNA-seq data with DESeq2. Genome Biol. 2014;15:550.

25 Salmon-Divon M, Dvinge H, Tammoja K, Bertone P. PeakAnalyzer: genome-wide annotation of chromatin binding and modification loci. BMC Bioinformatics. 2010;11:415.

26 Du Z, Zhou X, Ling Y, Zhang Z, Su Z. agriGO: a GO analysis toolkit for the agricultural community. Nucleic Acids Res. 2010;38:W64–70.

27 Tian T, Liu Y, Yan H, You Q, Yi X, Du Z, et al. agriGO v2.0: a GO analysis toolkit for the agricultural community, 2017 update. Nucleic Acids Res. 2017.

28 Machanick P, Bailey TL. MEME-ChIP: motif analysis of large DNA datasets. Bioinformatics. 2011;27:1696–7.

29 You Y, Sawikowska A, Neumann M, Pose D, Capovilla G, Langenecker T, et al. Temporal dynamics of gene expression and histone marks at the Arabidopsis shoot meristem during flowering.Nat Commun. 2017;8:15120.

30 Endo M, Shimizu H, Nohales MA, Araki T, Kay SA. Tissue-specific clocks in Arabidopsis show asymmetric coupling. Nature. 2014;515:419–22.

31 Szklarczyk D, Morris JH, Cook H, Kuhn M, Wyder S, Simonovic M, et al. The STRING database in 2017: quality-controlled protein-protein association networks, made broadly accessible. Nucleic Acids Res. 2017;45:D362–d8.

32 Cui D, Zhao J, Jing Y, Fan M, Liu J, Wang Z, et al. The arabidopsis IDD14, IDD15, and IDD16 cooperatively regulate lateral organ morphogenesis and gravitropism by promoting auxin biosynthesis and transport. PLoS Genet. 2013;9:e1003759.

33 Yoshida H, Hirano K, Sato T, Mitsuda N, Nomoto M, Maeo K, et al. DELLA protein functions as a transcriptional activator through the DNA binding of the indeterminate domain family proteins. Proc Natl Acad Sci U S A. 2014;111:7861–6.

34 Moreno-Risueno MA, Sozzani R, Yardimci GG, Petricka JJ, Vernoux T, Blilou I, et al. Transcriptional control of tissue formation throughout root development. Science. 2015;350:426–30.

35 Behringer C, Schwechheimer C. B-GATA transcription factors – insights into their structure, regulation, and role in plant development. Front Plant Sci. 2015;6:90.

36 De Rybel B, Vassileva V, Parizot B, Demeulenaere M, Grunewald W, Audenaert D, et al. A novel aux/IAA28 signaling cascade activates GATA23-dependent specification of lateral root founder cell identity. Curr Biol. 2010;20:1697–706.

37 Richter R, Behringer C, Zourelidou M, Schwechheimer C. Convergence of auxin and gibberellin signaling on the regulation of the GATA transcription factors GNC and GNL in Arabidopsis thaliana. Proc Natl Acad Sci U S A. 2013;110:13192–7.

38 Ranftl QL, Bastakis E, Klermund C, Schwechheimer C. LLM-Domain Containing B-GATA Factors Control Different Aspects of Cytokinin-Regulated Development in Arabidopsis thaliana. Plant Physiol. 2016;170:2295–311.

39 Grant CE, Bailey TL, Noble WS. FIMO: scanning for occurrences of a given motif. Bioinformatics. 2011;27:1017–8.

40 Chen H, Lai Z, Shi J, Xiao Y, Chen Z, Xu X. Roles of arabidopsis WRKY18, WRKY40 and WRKY60 transcription factors in plant responses to abscisic acid and abiotic stress. BMC Plant Biol. 2010;10:281.

41 Chen J, Nolan TM, Ye H, Zhang M, Tong H, Xin P, et al. Arabidopsis WRKY46, WRKY54, and WRKY70 Transcription Factors Are Involved in Brassinosteroid-Regulated Plant Growth and Drought Responses. Plant Cell. 2017;29:1425–39.

42 Birkenbihl RP, Kracher B, Somssich IE. Induced Genome-Wide Binding of Three Arabidopsis WRKY Transcription Factors during Early MAMP-Triggered Immunity. Plant Cell. 2017;29:20–38.

43 Yang Z, Tian L, Latoszek-Green M, Brown D, Wu K. Arabidopsis ERF4 is a transcriptional repressor capable of modulating ethylene and abscisic acid responses. Plant Mol Biol. 2005;58:585–96.

44 Son GH, Wan J, Kim HJ, Nguyen XC, Chung WS, Hong JC, et al. Ethylene-responsive element-binding factor 5, ERF5, is involved in chitin-induced innate immunity response. Mol Plant Microbe Interact. 2012;25:48–60.

45 Scarpeci TE, Frea VS, Zanor MI, Valle EM. Overexpression of AtERF019 delays plant growth and senescence, and improves drought tolerance in Arabidopsis. J Exp Bot. 2017;68:673–85.

46 Bolt S, Zuther E, Zintl S, Hincha DK, Schmulling T. ERF105 is a transcription factor gene of Arabidopsis thaliana required for freezing tolerance and cold acclimation. Plant Cell Environ. 2017;40:108–20.

47 Koyama T, Furutani M, Tasaka M, Ohme-Takagi M. TCP transcription factors control the morphology of shoot lateral organs via negative regulation of the expression of boundary-specific genes in Arabidopsis. Plant Cell. 2007;19:473–84.

48 Li S. The Arabidopsis thaliana TCP transcription factors: A broadening horizon beyond development. Plant Signal Behav. 2015;10:e1044192.

49 Alvarez JP, Furumizu C, Efroni I, Eshed Y, Bowman JL. Active suppression of a leaf meristem orchestrates determinate leaf growth. Elife. 2016;5.

50 Danisman S. TCP Transcription Factors at the Interface between Environmental Challenges and the Plant’s Growth Responses. Front Plant Sci. 2016;7:1930.

51 Stephenson PG, Fankhauser C, Terry MJ. PIF3 is a repressor of chloroplast development. Proc Natl Acad Sci U S A. 2009;106:7654–9.

52 Leivar P, Quail PH. PIFs: pivotal components in a cellular signaling hub. Trends Plant Sci. 2011;16:19–28.

53 Zhang Y, Mayba O, Pfeiffer A, Shi H, Tepperman JM, Speed TP, et al. A quartet of PIF bHLH factors provides a transcriptionally centered signaling hub that regulates seedling morphogenesis through differential expression-patterning of shared target genes in Arabidopsis. PLoS Genet. 2013;9:e1003244.

54 Pfeiffer A, Shi H, Tepperman JM, Zhang Y, Quail PH. Combinatorial complexity in a transcriptionally centered signaling hub in Arabidopsis. Mol Plant. 2014;7:1598–618.

55 Saini S, Sharma I, Pati PK. Versatile roles of brassinosteroid in plants in the context of its homoeostasis, signaling and crosstalks. Front Plant Sci. 2015;6:950.

56 Singh AP, Savaldi-Goldstein S. Growth control: brassinosteroid activity gets context. J Exp Bot. 2015;66:1123–32.

57 Yu X, Li L, Zola J, Aluru M, Ye H, Foudree A, et al. A brassinosteroid transcriptional network revealed by genome-wide identification of BESI target genes in Arabidopsis thaliana. Plant J. 2011;65:634–46.

58 Chen D, Xu G, Tang W, Jing Y, Ji Q, Fei Z, et al. Antagonistic basic helix-loop-helix/bZIP transcription factors form transcriptional modules that integrate light and reactive oxygen species signaling in Arabidopsis. Plant Cell. 2013;25:1657–73.

59 Toledo-Ortiz G, Johansson H, Lee KP, Bou-Torrent J, Stewart K, Steel G, et al. The HY 5-PIF regulatory module coordinates light and temperature control of photosynthetic gene transcription. PLoS Genet. 2014;10:e1004416.

60 Neph S, Stergachis AB, Reynolds A, Sandstrom R, Borenstein E, Stamatoyannopoulos JA. Circuitry and dynamics of human transcription factor regulatory networks. Cell. 2012;150:1274–86.

61 Murashige T, Skoog F. A Revised Medium for Rapid Growth and Bioassays with Tobacco Tisue Cultures. Physiol Plantarum. 1962;15:473–97.

62 Clough SJ, Bent AF. Floral dip: a simplified method for Agrobacterium-mediated transformation of Arabidopsis thaliana. Plant J. 1998;16:735–43.

63 Brand U, Grunewald M, Hobe M, Simon R. Regulation of CLV3 expression by two homeobox genes in Arabidopsis. Plant Physiol. 2002;129:565–75.

64 Zilberman D, Coleman-Derr D, Ballinger T, Henikoff S. Histone H2A.Z and DNA methylation are mutually antagonistic chromatin marks. Nature. 2008;456:125–9.

65 Langmead B, Salzberg SL. Fast gapped-read alignment with Bowtie 2. Nat Methods. 2012;9:357–9.

66 Li H, Handsaker B, Wysoker A, Fennell T, Ruan J, Homer N, et al. The Sequence Alignment/Map format and SAMtools. Bioinformatics. 2009;25:2078–9.

67 Thorvaldsdottir H, Robinson JT, Mesirov JP. Integrative Genomics Viewer (IGV): high-performance genomics data visualization and exploration. Brief Bioinform. 2013;14:178–92.

68 Huang W, Loganantharaj R, Schroeder B, Fargo D, Li L. PAVIS: a tool for Peak Annotation and Visualization. Bioinformatics. 2013;29:3097–9.

69 Anders S, Pyl PT, Huber W. HTSeq-- a Python framework to work with high-throughput sequencing data. Bioinformatics. 2015;31:166–9.

70 Medina-Rivera A, Defrance M, Sand O, Herrmann C, Castro-Mondragon JA, Delerce J, et al. RSAT 2015: Regulatory Sequence Analysis Tools. Nucleic Acids Res. 2015;43:W50–6.

71 Weirauch MT, Yang A, Albu M, Cote A, Montenegro-Montero A, Drewe P, et al. Determination and inference of eukaryotic transcription factor sequence specificity. Cell. 2014;158:1431–43.

72 O'Malley RC, Huang SS, Song L, Lewsey MG, Bartlett A, Nery JR, et al. Cistrome and Epicistrome Features Shape the Regulatory DNA Landscape. Cell. 2016;165:1280–92.

73 Pedmale UV, Huang SS, Zander M, Cole BJ, Hetzel J, Ljung K, et al. Cryptochromes Interact Directly with PIFs to Control Plant Growth in Limiting Blue Light. Cell. 2016;164:233–45.

